# Gene model for the ortholog of *Lst8* in *Drosophila yakuba*

**DOI:** 10.64898/2026.05.12.723325

**Authors:** Megan E. Lawson, Kylee Sanow, Kuntepi Chetana, Elizabeth Taylor, Ashley Morgan, Devan Flannery, Courtney Elsie, Chinmay P. Rele, Laura K. Reed, Kellie S. O’Rourke

## Abstract

Gene model for the ortholog of *Lst8* (*Lst8*) in the May 2011 (WUGSC dyak_caf1/DyakCAF1) Genome Assembly (GenBank Accession: GCA_000005975.1) of *Drosophila yakuba*. This ortholog was characterized as part of a developing dataset to study the evolution of the Insulin/insulin-like growth factor signaling pathway (IIS) across the genus *Drosophila* using the Genomics Education Partnership gene annotation protocol for Course-based Undergraduate Research Experiences.

## Introduction

“Computational gene predictions in non-model organisms often can be improved by careful manual annotation and curation, allowing for more accurate analyses of gene and genome evolution (Mudge and Harrow 2016; Tello-Ruiz et al., 2019). The Genomics Education Partnership (thegep.org) uses web-based tools to allow undergraduates to participate in course-based research by generating manual annotations of genes in non-model species (Rele et al., 2023). These models of orthologous genes across species, such as the one presented here, then provide a reliable basis for further evolutionary genomic analyses when made available to the scientific community. The particular gene ortholog described here *Lst8* (*Lst8*) in *D. yakuba* was characterized as part of a developing dataset to study the evolution of the Insulin/insulin-like growth factor signaling pathway (IIS) across the genus *Drosophila*.” (Myers et al., 2024).

“*D. yakuba* (Taxonomic ID: 7245) is part of the *melanogaster* species group within the subgenus *Sophophora* of the genus *Drosophila* (Sturtevant 1939; Bock and Wheeler 1972). It was first described by Burla (1954). *D. yakuba* is widespread in sub-Saharan Africa and Madagascar (Lemeunier et al., 1986; https://www.taxodros.uzh.ch, accessed 1 Feb 2023; Markow and O’Grady 2006) where figs served as a primary host along with other rotting fruits (Lachaise and Tsacas 1983).” (Koehler et al., 2024).

“The IIS pathway is a highly conserved signaling pathway in animals and is central to mediating organismal responses to nutrients (Hietakangas and Cohen 2009; Grewal 2009)” (Myers et al., 2024). “The *Lst8* gene was first identified in *Saccharomyces cerevisiae* and was shown to be a component of the target-of-rapamycin (TOR) protein complexes (Roberg et al., 1997; Loewith et al., 2002). In *Drosophila, Lst8* (FBgn0264691) was cloned as a component of the TORC1 and TORC2 protein complexes and was shown to be essential for TORC2 function, but not for TORC1 function (Wang et al., 2012). While an RNAi knockdown of *Lst8* does not show an obvious phenotype (Yang et al., 2006), a *Lst8* knockout mutant shows growth defects consistent with the role of TOR signaling in nutrient sensing and growth (Wang et al, 2012). TORC2 regulates cell growth via a Myc-dependent pathway, and growth defects in the *Lst8* knockout mutant can be rescued by Myc expression (Kuo et al., 2015). The human ortholog mLST8 is also a subunit of mTORC1 and mTORC2 and increased expression of mLST8 has been shown to play a role in tumor progression (Kakumoto et al., 2015).” (Myers et al., 2026)

The model presented here is the ortholog of *Lst8* in the May 2011 (WUGSC dyak_caf1/DyakCAF1) assembly of *D. yakuba* (GCA_000005975.1) and corresponds to the Gnomon Peptide ID XP_002101753.1 predicted model in *D. yakuba* (LOC6525933). This gene model is based on RNA-Seq data from *D. yakuba* (Gravely *et al*., 2011; SRP006203) and the *Lst8* (GCA_000001215.4 – Drosophila 12 Genomes Consortium *et al*., 2007) in *D. melanogaster* from FB2022_03 (Larkin *et al*., 2021). The complete methods and dataset versions used to establish the gene model are described in Rele *et al*. (2020). The Genomics Education Partnership maintains a mirror of the UCSC Genome Browser (Kent WJ et al., 2002; Gonzalez et al., 2021), which is available at https://gander.wustl.edu.

## Results

### Synteny

*Lst8* occurs on chromosome X in *D. melanogaster* and is flanked upstream by *Hecw* and *LysRS* and downstream by *Gga* and *Hex-A*. We determined that the putative ortholog of *Lst8* is found on chromosome X (CM000162.2) in *D. yakuba* with LOC6525933 (XP_002101753.1) (via *blastp* search with an e-value of 0.0 and percent identity of 98.08%). It is flanked upstream by LOC6525935 and LOC6525934, which correspond to *Hecw* and *LysRS* in *D. melanogaster* with e-values of 0.0 and 0.0 respectively and percent identities of 93.35% and 95.62% respectively, as determined by *blastp*. It is flanked downstream by LOC6525932 and LOC6525931, which correspond to *Gga* and *Hex-A* in *D. melanogaster* with e-values of 0.0 and 0.0 respectively and percent identities of 89.07% and 96.15% respectively, as determined by *blastp* (**Figure 1A**, Altschul et al., 1990). We believe this is the correct ortholog assignment for *Lst8* in *D. yakuba* because the local synteny of the genomic neighborhood is completely conserved between the two species, and the *blastp* matches used to determine orthology are all very high quality.

**Figure 1:**
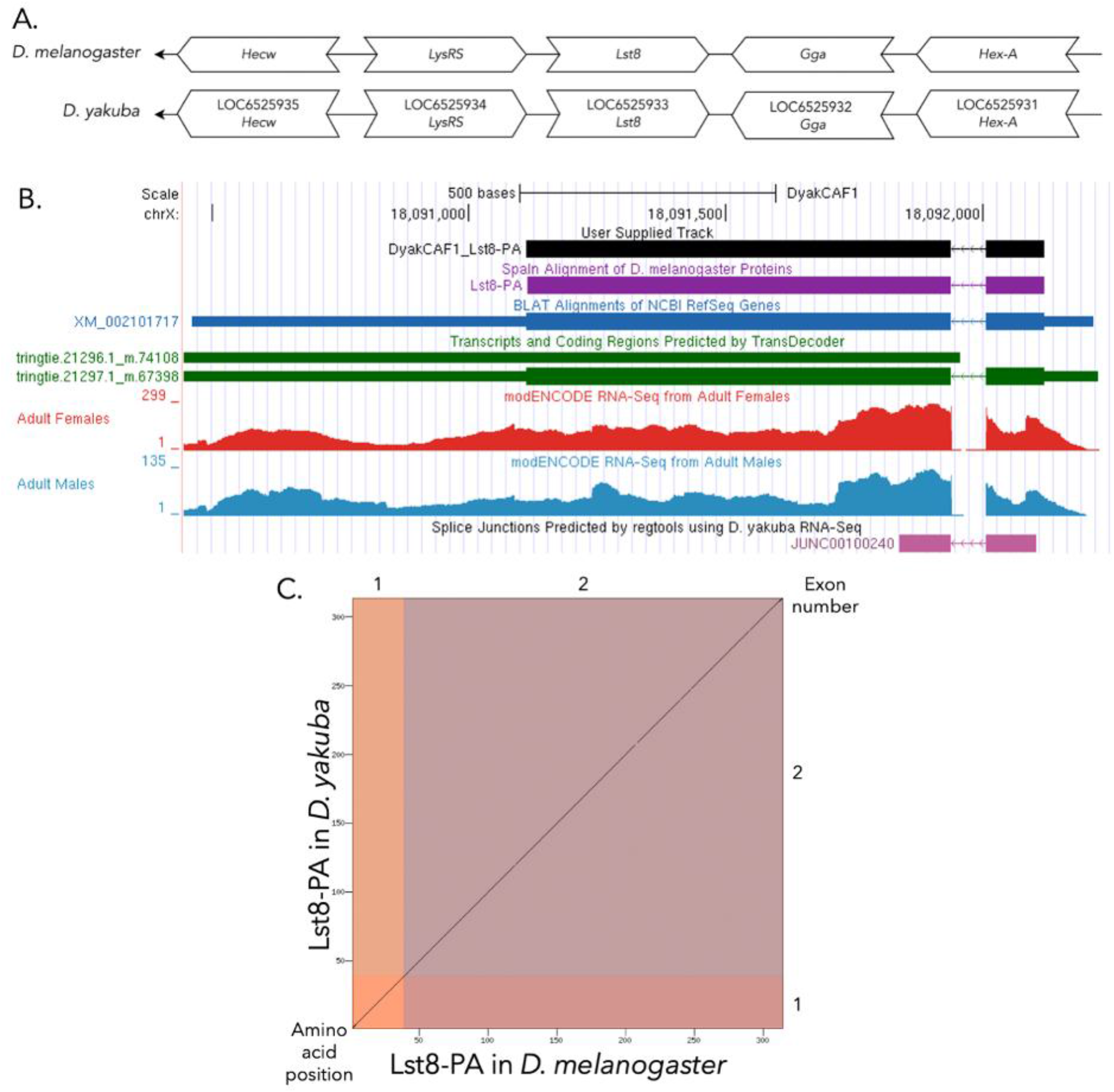
(A) **Synteny of genomic neighborhood of *Lst8* in *D. melanogaster* and *D. yakuba***. Gene arrows pointing in the same direction as target gene in both *D. yakuba* and *D. melanogaster* are on the same strand as the target gene; gene arrows pointing in the opposite direction are on the opposite strand. The thin underlying arrows pointing to the left indicate that *Lst8* is on the negative strand. White arrows in *D. yakuba* indicate the locus ID and the orthology to the corresponding gene in *D. melanogaster*. The gene names given in the *D. yakuba* gene arrows indicate the orthologous gene in *D. melanogaster*, while the locus identifiers are specific to *D. yakuba*. (B) **Gene Model in UCSC Track Hub** (Raney et al. 2014): the gene model in *D. yakuba* (black), Spaln of D. melanogaster Proteins (purple, alignment of RefSeq proteins from *D. melanogaster*), BLAT alignments of NCBI RefSeq Genes (blue, alignment of RefSeq genes for *D. yakuba*), RNA-Seq from adult females (red) and adult males (blue) (alignment of Illumina RNA-Seq reads from *D. yakuba*), and Transcripts (green) including coding regions predicted by TransDecoder and Splice Junctions Predicted by regtools using *D. yakuba* RNA-Seq (Gravely *et al*., 2011; SRP006203). The splice junction shown has a read-depth of 317. The custom gene model (User Supplied Track) is indicated in black with exons depicted by boxes and introns by narrow lines (arrows indicate direction of transcription). (C) **Dot Plot of Lst8-PA in *D. melanogaster* (*x*-axis) vs. the orthologous peptide in *D. yakuba* (*y*-axis)**. Amino acid number is indicated along the left and bottom; exon number is indicated along the top and right, and exons are also highlighted with alternating colors.

### Protein Model

*Lst8* in *D. yakuba* has one protein-coding isoform, Lst8-PA (**Figure 1B**). Isoform Lst8-PA contains two protein coding exons. This is the same relative to the ortholog in *D. melanogaster*, which also has one protein-coding isoform with two exons. The sequence of *Lst8* in *D. yakuba* has 98.08% identity with the *Lst8* in *D. melanogaster* as determined by *blastp* (**Figure 1C**).

## Methods

Detailed methods including algorithms, database versions, and citations for the complete annotation process can be found in Rele *et al*. (2020).

## Acknowledgements

We would like to thank Wilson Leung for developing and maintaining the technological infrastructure that was used to create this gene model and Laura K. Reed for overseeing the project.

## Funding

This material is based upon work supported by the National Science Foundation (1915544) and the National Institute of General Medical Sciences of the National Institutes of Health (R25GM130517) to the Genomics Education Partnership (GEP; https://thegep.org/; PI-LKR). Any opinions, findings, and conclusions or recommendations expressed in this material are solely those of the author(s) and do not necessarily reflect the official views of the National Science Foundation nor the National Institutes of Health.

## Supplemental Files

1. Zip file containing a FASTA, PEP, GFF files for the gene model

## Metadata

Bioinformatics, Genomics, *Drosophila*, Genotype Data, New Finding

## Notes

### Competing Interest Statement

The authors have declared no competing interest.

## References

Altschul, S. F., Gish, W., Miller, W., Myers, E. W., & Lipman, D. J. (1990). Basic local alignment search tool. Journal of Molecular Biology, 215(3), 403–410. 10.1016/S0022-2836(05)80360-2

Bächli, G. (2026, February 4). TaxoDros. TaxoDros. https://www.taxodros.uzh.ch/

Bock IR, Wheeler MK. 1972. The Drosophila melanogaster species group. University of Texas Publications, 7213, 1–102.

Burla, H. (1954) Zur Kenntnis der Drosophiliden der Elfenbeinkuste (Franzosisch West-Afrika). Revue Suisse de Zoologie, 61, (Suppl.), 1–218.

Drosophila 12 Genomes Consortium. (2007). Evolution of genes and genomes on the Drosophila phylogeny. Nature, 450(7167), 203–218. 10.1038/nature06341

Gonzalez, J. N., Zweig, A. S., Speir, M. L., Schmelter, D., Rosenbloom, K. R., Raney, B. J., Powell, C. C., Nassar, L. R., Maulding, N. D., Lee, C. M., Lee, B. T., Hinrichs, A. S., Fyfe, A. C., Fernandes, J. D., Diekhans, M., Clawson, H., Casper, J., Benet-Pagès, A., Barber, G. P., … Kent, W. J. (2021). The UCSC Genome Browser database: 2021 update. Nucleic Acids Research, 49(D1), D1046–D1057. 10.1093/nar/gkaa1070

Gravely, B. R., Brooks, A. N., Carlson, J. W., Duff, M. O., Landolin, J. M., Yang, L., Artieri, C. G., Van Baren, M. J., Boley, N., Booth, B. W., Brown, J. B., Cherbas, L., Davis, C. A., Dobin, A., Li, R., Lin, W., Malone, J. H., Mattiuzzo, N. R., Miller, D., … Celniker, S. E. (2011). The developmental transcriptome of Drosophila melanogaster. Nature, 471(7339), 473–479. 10.1038/nature09715

Grewal, S. S. (2009). Insulin/TOR signaling in growth and homeostasis: A view from the fly world. The International Journal of Biochemistry & Cell Biology, 41(5), 1006–1010. 10.1016/j.biocel.2008.10.010

Hietakangas, V., & Cohen, S. M. (2009). Regulation of Tissue Growth through Nutrient Sensing. Annual Review of Genetics, 43(1), 389–410. 10.1146/annurev-genet-102108-134815

Kakumoto, K., Ikeda, J., Okada, M., Morii, E., & Oneyama, C. (2015). mLST8 Promotes mTOR-Mediated Tumor Progression. PLOS ONE, 10(4), e0119015. 10.1371/journal.pone.0119015

Kent, W. J., Sugnet, C. W., Furey, T. S., Roskin, K. M., Pringle, T. H., Zahler, A. M., & Haussler, A. D. (2002). The Human Genome Browser at UCSC. Genome Research, 12(6), 996–1006. 10.1101/gr.229102

Koehler AC, Romo I, Le V, Romo I, Youngblom JJ, Hark AT, Rele CP. 2024. Gene model for the ortholog of GlyS in Drosophila yakuba. microPublication Biology. (submitted)

Kuo, Y., Huang, H., Cai, T., & Wang, T. (2015). Target of Rapamycin Complex 2 regulates cell growth via Myc in Drosophila. Scientific Reports, 5(1), 10339. 10.1038/srep10339

Larkin, A., Marygold, S. J., Antonazzo, G., Attrill, H., dos Santos, G., Garapati, P. V., Goodman, J. L., Gramates, L. S., Millburn, G., Strelets, V. B., Tabone, C. J., Thurmond, J., FlyBase Consortium, Perrimon, N., Gelbart, S. R., Agapite, J., Broll, K., Crosby, M., Dos Santos, G., … Lovato, T. (2021). FlyBase: Updates to the Drosophila melanogaster knowledge base. Nucleic Acids Research, 49(D1), D899–D907. 10.1093/nar/gkaa1026

Lachaise D, Tsacas L. 1983. Breeding-sites of tropical African Drosophilids. Ashburner, Carson, Thompson, 1981-1986. 3d: 21.

Lemeunier, F., David, J., Tsacas, L., & Ashburner, M. (1986). The melanogaster species group. In M. Ashburner, J. Thompson, & H. Carson (Eds.), The Genetics and Biology of Drosophila (Vol. 3e, pp. 147–256). Academic Press.

Loewith, R., Jacinto, E., Wullschleger, S., Lorberg, A., Crespo, J. L., Bonenfant, D., Oppliger, W., Jenoe, P., & Hall, M. N. (2002). Two TOR Complexes, Only One of which Is Rapamycin Sensitive, Have Distinct Roles in Cell Growth Control. Molecular Cell, 10(3), 457–468. 10.1016/S1097-2765(02)00636-6

Markow, T. A., & O’Grady, P. M. (2006). Drosophila.

Mudge, J. M., & Harrow, J. (2016). The state of play in higher eukaryote gene annotation. Nature Reviews Genetics, 17(12), 758–772. 10.1038/nrg.2016.119

Myers A, Hoffman A, Natysin M, Arsham AM, Stamm J, Thompson JS, Rele CP, Reed LK. 2024. Gene model for the ortholog of Myc in Drosophila ananassae, micropublication Biology.

Myers A.R., Wellik I.G., Duan B., Herrenkohl E., Yu A.M., Agrimson K.S., Sustacek M.K., Yang M., Thompson J.S., Rele C.P. 2024. Gene model for the ortholog of Lst8 in Drosophila grimshawi, micropublication Biology (submitted)

Raney, B. J., Dreszer, T. R., Barber, G. P., Clawson, H., Fujita, P. A., Wang, T., Nguyen, N., Paten, B., Zweig, A. S., Karolchik, D., & Kent, W. J. (2014). Track data hubs enable visualization of user-defined genome-wide annotations on the UCSC Genome Browser. Bioinformatics, 30(7), 1003–1005. 10.1093/bioinformatics/btt637

Rele CP, Sandlin KM, Leung W, Reed LK. 2020. Manual Annotation of Genes within Drosophila Species: the Genomics Education Partnership protocol. bioRxiv 2020.12.10.420521 doi: 10.1101/2020.12.12.420521

Rele, C. P., Sandlin, K. M., Leung, W., & Reed, L. K. (2023). Manual annotation of Drosophila genes: A Genomics Education Partnership protocol. F1000Research, 11, 1579. 10.12688/f1000research.126839.3

Roberg KJ, Bickel S, Rowley N, Kaiser CA. 1997. Control of amino acid permease sorting in the late secretory pathway of Saccharomyces cerevisiae by SEC13, LST4, LST7 and LST8. Genetics 147:1569–1584

Sturtevant, A. H. (1939). On the Subdivision of the Genus Drosophila. Proceedings of the National Academy of Sciences, 25(3), 137–141. 10.1073/pnas.25.3.137

Tello-Ruiz, M. K., Marco, C. F., Hsu, F.-M., Khangura, R. S., Qiao, P., Sapkota, S., Stitzer, M. C., Wasikowski, R., Wu, H., Zhan, J., Chougule, K., Barone, L. C., Ghiban, C., Muna, D., Olson, A. C., Wang, L., Ware, D., & Micklos, D. A. (2019). Double triage to identify poorly annotated genes in maize: The missing link in community curation. PLOS ONE, 14(10), e0224086. 10.1371/journal.pone.0224086

Wang, T., Blumhagen, R., Lao, U., Kuo, Y., & Edgar, B. A. (2012). LST8 Regulates Cell Growth via Target-of-Rapamycin Complex 2 (TORC2). Molecular and Cellular Biology, 32(12), 2203–2213. 10.1128/MCB.06474-11

Yang, Q., Inoki, K., Ikenoue, T., & Guan, K.-L. (2006). Identification of Sin1 as an essential TORC2 component required for complex formation and kinase activity. Genes & Development, 20(20), 2820–2832. 10.1101/gad.1461206

